# Dectin-1 ligands produce distinct training phenotypes in human monocytes through differential activation of signaling networks

**DOI:** 10.1101/2023.10.01.560380

**Authors:** Quen J. Cheng, Kylie Farrell, Jeffrey Fenn, Zuchao Ma, Sara K. Makanani, Jonathan Siemsen

**Affiliations:** Department of Medicine, Division of Infectious Diseases, David Geffen School of Medicine, University of California, Los Angeles, CA, USA; Molecular Biology Institute, University of California, Los Angeles, California, USA; Department of Microbiology, Immunology, and Molecular Genetics, University of California, Los Angeles, CA, USA; Department of Surgery, Center of Excellence in Inflammation, Infectious Disease and Immunity, Quillen College of Medicine, East Tennessee State University, Johnson City, TN, USA

**Keywords:** Trained immunity, Innate immune memory, β-glucan, Dectin-1, toll-like receptors

## Abstract

Cells of the innate immune system retain memory of prior exposures through a process known as innate immune training. β-glucan, a Dectin-1 ligand purified from the *Candida albicans* cell wall, has been one of the most widely u:lized and well-characterized ligands for inducing innate immune memory. However, many Dectin-1 agonists exist, and it is not known whether all Dectin-1 ligands produce the same phenotype. Using a well-established *in vitro* model of trained immunity, we compared two commercially available Dectin-1 agonists, zymosan and depleted zymosan, with the gold standard β-glucan in the literature. We found that depleted zymosan, a β-glucan purified from *Saccharomyces cerevisiae* cell wall through alkali treatment, produced near identical training effects as *C. albicans* β-glucan. However, untreated zymosan produced a distinct training effect from β-glucans at both the transcript and cytokine level. Training with zymosan diminished, rather than potentiated, induction of key cytokines such as TNF, IL-12, and IL-6. Zymosan activated NF𝓀B and AP-1 transcription factors more strongly than β-glucans. The addition of the toll-like receptor (TLR) ligand Pam3CSK4 was sufficient to convert the training effect of β-glucans to a phenotype resembling training with zymosan. We conclude that differential activation of TLR signaling pathways determines the phenotype of innate immune training induced by Dectin-1. These findings bring clarity to the specific question of which Dectin-1 agonists produce prototypical training effects and provide broader insight into how signaling networks regulate innate immune training.

## Introduction

There is growing recognition that cells of the innate immune system, such as monocytes and macrophages, retain memory of prior exposures. This innate immune “training” produces phenotypic shifts that alter macrophage responses to subsequent immune threats (1,2). Innate immune training carries therapeutic potential to enhance immune responses to heterologous pathogens (3,4) or as a novel strategy for cancer immune therapy (5). Innate immune training may also contribute to the pathogenesis of both hyper- and hypo-inflammatory disease states (6,7).

Unlike T- and B-cells of the adaptive immune system, which encode memory of prior exposures through genetic rearrangement, innate immune cells retain memory through non- genetic mechanisms such as metabolic reprogramming (8) and epigenetic modifications (9). Importantly, while the most widely used training ligands increase (or “potentiate”) inflammatory cytokine production, other training ligands have been shown to diminish inflammatory cytokine production (10,11). A key question in the field is what determines whether a given training ligand and the cell signaling pathways it activates will produce potentiated or diminished inflammatory responses.

One archetypal training ligand has been the fungal pathogen, *Candida albicans*, and the primary pathogen-associated molecular pattern (PAMP) found in its cell wall, (1,3)-β-D-glucan (12–14). *In vivo* injection of β-glucan protects mice against subsequent infection with a broad range of heterologous pathogens (3). β-glucan exposure *in vivo* produces trained macrophages that respond to secondary lipopolysaccharide (LPS) s:mula:on with increased production of inflammatory cytokines (12). This potentiation of cytokine production is also observed in human monocyte-derived macrophages trained *in vitro* with β-glucan (15). The β-glucan training effect is dependent on the Dectin-1/Raf-1/mTOR signaling axis (12,16) and is correlated with changes in chromatin accessibility and H3K27ac marks (9) as well as a shih towards aerobic glycolysis (8).

Notably, these insights were gained from studies that utilized a preparation of β-glucan that is not commercially available but has been generously shared by the Williams lab at East Tennessee State University (ETSU). This highly purified (1→3,1→6)-β-glucan is isolated from *Candida albicans* SC5314 yeast cell walls and has been thoroughly characterized (17,18).

However, commercially available “β-glucan” products are much more chemically and structurally diverse. Zymosan, one of the most widely used PAMPs in fungal pathogenesis studies, is a complex mixture of unpurified *Saccharomyces cerevisiae* cell wall. Although β-glucan is the most prevalent macromolecule in zymosan, these particles also contain mannoproteins, lipids, protein, chitin, and other macromolecules in variable proportions (19) and therefore binds toll-like receptors (TLRs) in addition to Dectin-1 (20–22). “Depleted zymosan” is a commercially available product in which zymosan is treated with hot alkali. This treatment depletes mannoproteins and chitin, resulting in a relatively pure β-glucan derived from *S. cerevisiae* that binds Dectin-1 with higher specificity (23). Whether zymosan and depleted zymosan produce similar innate immune training phenotypes to the archetypal β-glucan from ETSU is not known.

We show here that different Dectin-1 ligands can in fact produce distinct phenotypes of innate immune training. The commercially available β-glucan, depleted zymosan, has nearly identical effects to ETSU β-glucan. But untreated zymosan produces a distinct training regimen, exerting the opposite effect on key inflammatory cytokines. These differences arise from zymosan-specific activation of NF𝓀B and AP-1 transcription factors and can be replicated by concurrent training with a TLR ligand plus β-glucan. Our study thus identifies important differences between fungal cell wall ligands and sheds light on underlying mechanisms that may determine the stimulus-specificity of innate immune training beyond these ligands.

## Methods

### Reagents

*C. albicans* β-glucan was generously provided by Dr. David Williams at ETSU. The identity and structure of ETSU β-glucan was characterized by NMR and confirmed to be >95% (1→3,1→6)-β-D-glucan. Sterility and endotoxin testing of the glucan was confirmed as previously described (24). Zymosan and depleted zymosan were purchased from InvivoGen and validated to be endotoxin-free by the vendor.

### Tissue Culture

Peripheral blood mononuclear cells were obtained from healthy donors through the UCLA-CFAR Virology Core. Monocytes were isolated by negative bead selection using a Pan-Monocyte Isolation Kit (Miltenyi Biotec 130-096-537) and grown on tissue culture plates (Corning) in complete medium consisting of RPMI 1640 broth containing 10% fetal bovine serum (Gibco), 100 IU/ml penicillin streptomycin, 100 μg/ml streptomycin, 2 mM L-glutamine, and 10 ng/ml recombinant human M-CSF (R&D Systems 216-MC-100) at 37°C for seven days.

### In vitro trained immunity

Monocytes were rested in complete medium for 30-60 minutes after purification, then trained with the following ligands for 24 hours: zymosan (1 μg/mL, InvivoGen tlrl-zyn), ETSU β-glucan (10 μg/mL), depleted zymosan (10 μg /mL, InvivoGen tlrl-zyd), Pam3CSK4 (100 ng/mL, Invivogen tlrl-pms), or IFN-β (10 U/mL, PBL Assay Science 11415-1). Aher 24 hours of training, the medium was aspirated, cells were washed with warm HBSS, and fresh complete medium was added. On Day 4, medium was aspirated and fresh medium was added. On Day 7, the cells were stimulated with LPS (1 ng/mL, Millipore Sigma L6529) and analyzed for cytokine release or gene expression.

### Cytokine release assays

For cytokine release assays, monocytes were plated at different concentrations to account for the variable effects of trained immunity on cell proliferation rates (15). On Day 7, wells with similar cell densities by microscopy were chosen for comparisons between training regimens. Medium was removed, centrifuged to remove non-adherent cells, and snap-frozen for cytokine measurement. TNF-α concentration was determined by sandwich ELISA (R&D Systems DY210-05) per manufacture protocol. Multiplex bead-based measurements of cytokines (Luminex, Thermo Fisher) were performed by the UCLA Immune Assessment Core.

### Luminex analysis

Data from a panel of 38 analytes were quantified using standard curves provided by the manufacturer. Analytes expressed at levels below the lowest standard were excluded from downstream analysis. To account for variability in cell density and absolute cytokine concentrations between biological replicates, each replicate was Z-scored across training conditions.

### RNA Extraction and RT-qPCR

Cells were lysed using Qiagen RNeasy Micro Kit (Qiagen 74004), and RNA was extracted according to manufacturer’s protocol. Reverse transcription was carried out using LunaScript RT (New England Biolabs E3010) per manufacturer’s protocol. Quantitative PCR was performed with Luna® Universal qPCR Master Mix (New England Biolabs M3003). The following primer sequences were used: *IL12B –* GCCCAGAGCAAGATGTGTCA, CACCATTTCTCCAGGGGCAT; and *HPRT* – AGGACTGAACGTCTTGCTCG, ATCCAACACTTCGTGGGGTC.

### RNA-sequencing

RNA-seq libraries were prepared using KAPA stranded mRNA-seq library kit per manufacturer’s instructions and single-end sequenced at read length 50 bp on an Illumina HiSeq 2500 to a depth of 15-30 million reads per library. The low quality 3’ends of reads were trimmed (cutoff q=30), and remaining adapter sequences were removed using cutadapt (25). Processed reads were aligned to hg38 genome using STAR (26), and count tables were generated by the featureCount function in deepTools (27). Counts were normalized using the TMM-normalization method and RPKM values were generated using edgeR (28). Genes below an expression threshold of 4 CPM in all samples were excluded from downstream analysis. LPS-inducible genes were identified by FDR<0.05 and log2(fold-change)>2 compared to unstimulated. Potentiated or diminished LPS-responses were identified by refitting the linear model to the 603 LPS-inducible genes and imposing thresholds of p-value<0.05 and log2(fold-change)>0.5 comparing trained *vs*. untrained gene expression. For log2 transformations and fold-change calculations a pseudocount of 0.63 RPKM was added. The data were visualized using pheatmap and ggplot2 packages (29,30).

### Nuclear magnetic resonance

Zymosan, depleted zymosan, and *C. albicans* β-glucan were dissolved in DMSO-d6 at 10 mg/ml and 20 μL TFA-d was added. NMR data was collected at 60°C on a Bruker AvanceCore 400 spectrometer with 65,536 data points, 2 dummy scans, 128 scans, 1-sec pulse delay and 20.5 ppm sweep width centered at 6.175 ppm. Identification of glucan, protein, and lipids was performed as previously described (24).

### Immunoblots

Monocytes were isolated from PBMCs as above and cultured for 24 hours in M-CSF-containing medium to allow for adherence. Cells were then stimulated with the same concentrations of ligands as in the trained immunity assays. At the designated timepoints, cells were rinsed with cold PBS and detached using a cell scraper. The cytoplasmic membrane was lysed in hypotonic buffer (10 mM 7.9 pH HEPES, 10 mM KCl, 0.1 mM EGTA, 0.1mM EDTA) containing 1% NP-40. Nuclei were pelleted by centrifugation, and nuclear proteins were extracted in hypertonic solution (20 mM 7.9 pH HEPES, 420 mM NaCl, 1.5 mM MgCl2, 0.2 mM EDTA, 25% Glycerol) with protease and phosphatase inhibitors.

Protein concentrations were measured using Bradford Protein Assay (BioRad) and normalized in nuclear lysis buffer. 2x Laemmli Buffer (BioRad) containing 5% beta-mercaptoethanol was added, and samples were heated at 95°C for 10-20 minutes, separated by SDS-PAGE, and transferred to nitro-cellulose membranes. The following antibodies were used for immunoblotting: p-cJUN (Rabbit IgG, 1:2000, Cell Signaling Technology 9164), pIRF-3 (Rabbit IgG, 1:1000, Cell Signaling Technology 4947), p65(RelA) (Rabbit IgG, 1:1000, Santa Cruz 372), p-STAT1 (Rabbit IgG, 1:1000, Cell Signaling Technology 9167), and p84 (Rabbit IgG, 1:10000, 5E10), and an:-rabbit IgG (Goat IgG-HRP, 1:5000, Cell Signaling Technology 7074). Immunoblots were developed using chemiluminescent substrate (Supersignal West Pico Plus, Thermo Fisher) and visualized using a ChemiDoc MP imaging system (Bio-Rad, Hercules, CA). Following imaging, western blots were stripped for re-probing using a stripping buffer (ThermoFisher Scientific).

## Results

### Training with zymosan and β-glucans induce different cytokine responses to LPS

To determine the specificity of different fungal cell wall ligands in trained immunity, we employed a well-established *in vitro* trained immunity model using primary human monocytes (31). Monocytes were isolated from healthy human donors and differentiated to macrophages with MCSF, with or without training on Day 0 **[Fig. 1A]**. On Day 7, we assessed response to secondary stimulation by measuring TNF production after stimulation with low-dose LPS (1 ng/mL). Consistent with the literature, TNF production was potentiated when macrophages were trained with *C. albicans* β-glucan (β-gluETSU) with a mean fold-change of 1.99 relative to Untrained (p = 0.002) **[Fig. 1B]**. Training with depleted zymosan (β-gluDZ) also potentiated TNF production, with mean fold-change of 1.97 (p < 0.001). Surprisingly, we found that training with untreated zymosan produced the opposite effect, diminishing TNF production with a mean fold-change of 0.34 (p < 0.001). This effect was consistent across a range of doses **[Fig. S1]**.

**Figure 1:**
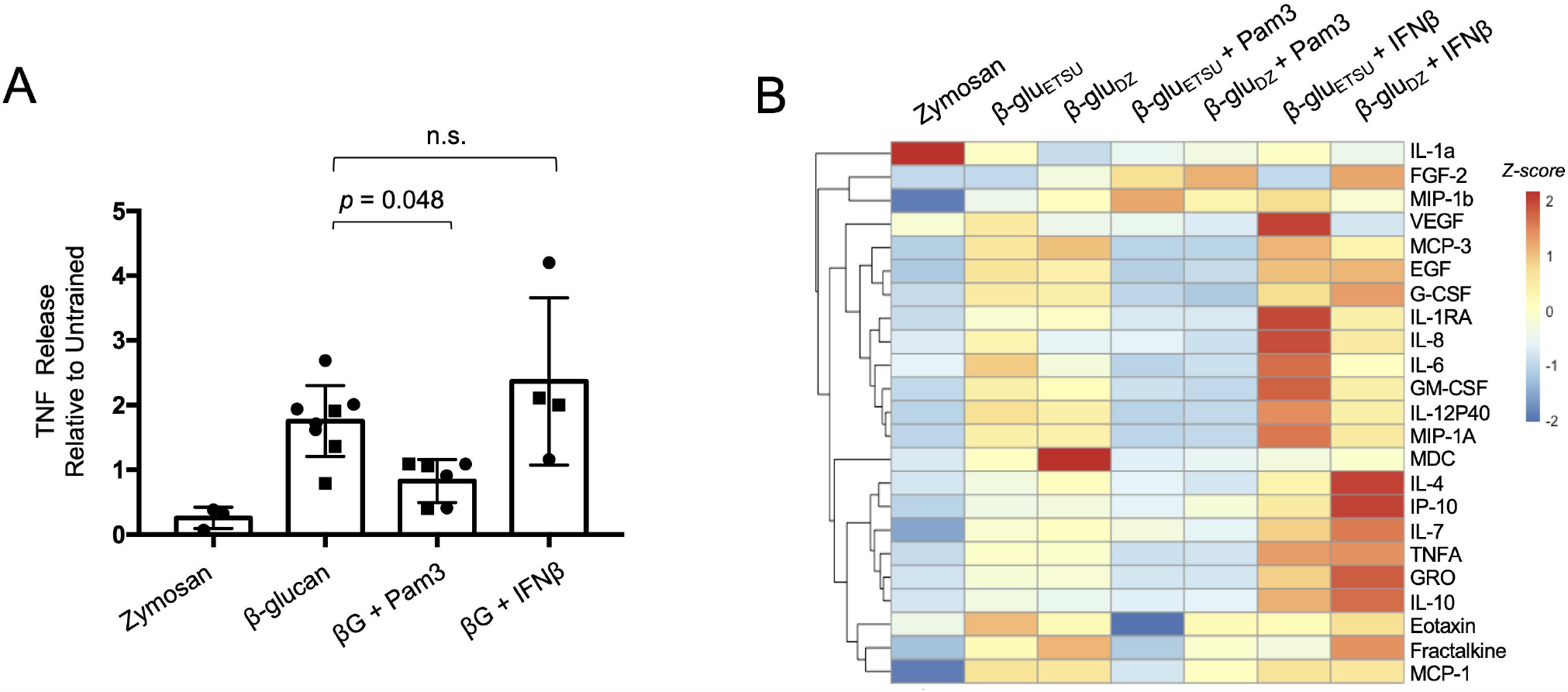
Training with zymosan and β-glucans induce different cytokine responses to LPS. **(A)** Experimental scheme. **(B)** TNF ELISA from supernatants of macrophages stimulated with LPS for eight hours, after training with the indicated ligands; p-values derived from two-tailed t-test; β-gluETSU = β-glucan from East Tennessee State University, β-gluDZ = depleted zymosan. **(C)** Heat map of multiplexed cytokine bead array from supernatant of macrophages stimulated with LPS for eight hours, after training with the indicated ligands; replicate columns within a condition are from a different donors.

To characterize training effects on a larger number of secreted proteins, we performed multiplexed bead-array cytokine measurements. Unsupervised hierarchical clustering revealed that a wide range of cytokines and chemokines behaved similarly to TNF, with diminished production in zymosan-trained macrophages and stable or potentiated production in depleted zymosan or ETSU β-glucan-trained macrophages **[Fig. 1C]**. These included classic inflammatory cytokines TNF, IL-6, and IL-12P40; chemokines critical for phagocyte recruitment such as IP-10, MIP-1a, and MIP-1b; and growth factors such as GM-CSF and G-CSF.

Interestingly, this effect was not universal, and individual cytokines displayed nuanced effects. For instance, IL-1α was potentiated by all three training regimens, especially zymosan, and IL-10 was more strongly potentiated by ETSU β-glucan than depleted zymosan. These data indicated that the effects of innate immune training are gene-specific, and that distinct regulatory strategies may exist for individual cytokines. Broadly speaking, however, we concluded that ETSU β-glucan and depleted zymosan produced similar training effects, in stark contrast to zymosan which diminished the production of the majority of cytokines.

### Training with zymosan and β-glucans induce different transcriptomic responses to LPS

Having established the differential training effects on secretion of cytokines, we wondered whether zymosan and β-glucans also exerted distinct effects on transcriptomic responses to LPS. We therefore trained monocytes and differentiated them to macrophages as before, then performed RNA-seq analysis. Samples were collected on Day 7 prior to secondary LPS stimulation (0 hour), and at one and three hours after LPS exposure. Experiments were performed in biological replicates with monocytes from different donors. We focused our analysis on 603 genes that were inducible by LPS in any training regimen, using a false discovery rate (FDR) cutoff of 0.05 and a log2 fold-change cutoff of 1.0 **[Table S1]**. Using unsupervised k-means clustering, we identified five distinct patterns of gene expression **[Fig. 2A]**. Cluster 1 genes had diminished LPS-responsiveness when trained with zymosan, but potentiated LPS-responsiveness when trained with either ETSU β-glucan or depleted zymosan. The differences between zymosan and β-glucans was statistically significant, with median log2(RPKM) differences of 0.769 (*p* = 10^-15^) and 0.915 (*p* < 10^-16^) for ETSU β-glucan and depleted zymosan, respectively, for Cluster 1 genes **[Fig. 2B]**. In contrast, Cluster 5 genes had the opposite pattern: potentiation when trained with zymosan and diminished responsiveness when trained with a β-glucan. These differences were also statistically significant, with median log2(RPKM) differences of -0.892 (*p* < 10^-16^) and -0.887 (*p* < 10^-16^) for ETSU β-glucan and depleted zymosan, respectively **[Fig. 2B]**. Cluster 2, 3, and 4 genes had diminished LPS-responsiveness in all three training regimens; these three clusters were distinguished from one another based on kinetics of gene induction.

**Figure 2:**
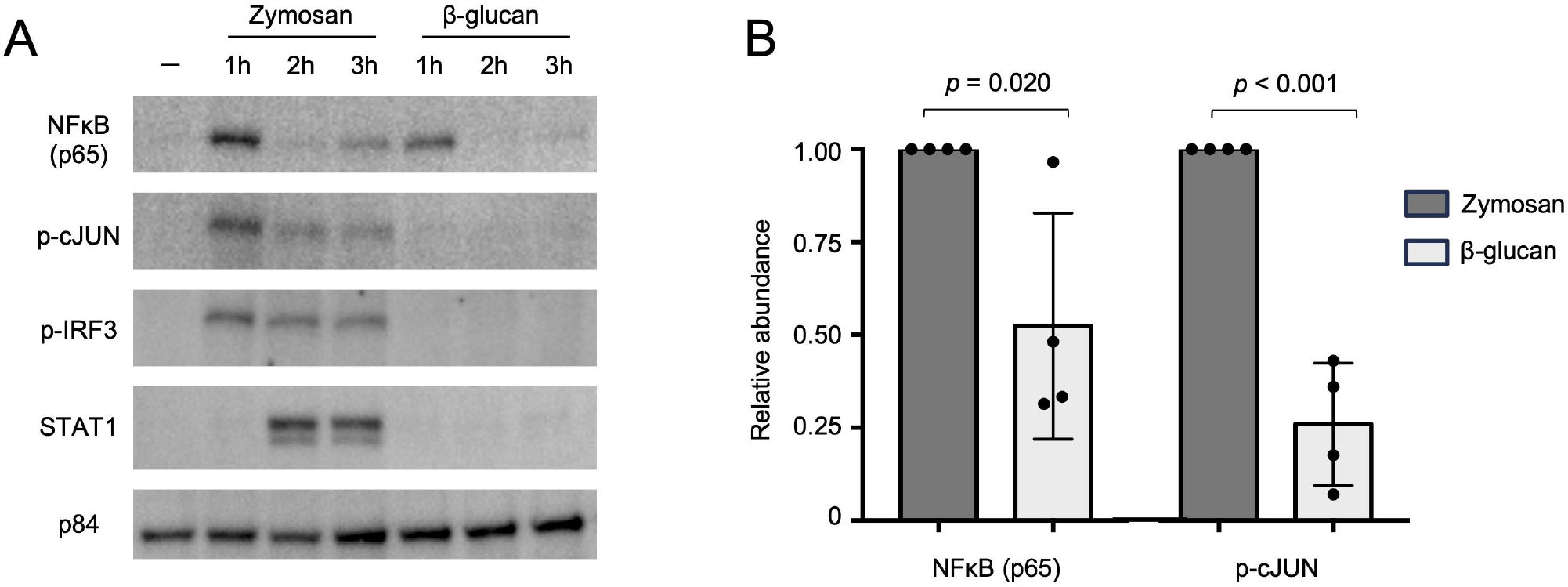
Training with zymosan and β-glucans induce different transcriptomic responses to LPS. **(A)** Heat map of 603 genes induced by LPS over a 3-hour time course in macrophages trained with the indicated ligands; average of two replicates; k-means clustering showing five distinct clusters. **(B)** Box and violin plots of RPKM at 3 hours for genes in Cluster 1 and Cluster 5; p-values from paired t-test. **(C)** Scatterplots comparing log fold-change of trained *vs*. untrained RPKM at 3-hour LPS timepoint; training effect of two types of β-glu are strongly correlated with each other (left), but weakly correlated with zymosan effects (right). **(D)** Heat map of correlation coefficients between conditions using the same analytical approach as (C). **(E)** Venn diagrams of the number of genes showing potentiated or diminished LPS response after training with Zymosan or either β-glucan, using statistical threshold of p < 0.05 and log fold-change > 0.5. **(F)** Venn diagram of genes with potentiated LPS response after training. **(G)** Expression of representative genes potentiated by Zymosan *(PTGIR, WNT5A)*, β-glucan *(IL12B*), or both *(IL1A)*; error bars = standard deviation, * = p < 0.05 in condition indicated by color.

Next, we quantified the differential effects of training. For each trained condition, we defined the training effect by calculating log2(fold-change) of trained *vs*. untrained at each timepoint. We found that training with ETSU β-glucan and depleted zymosan produced highly correlated effects; for example, at the 3-hour timepoint, the correlation coefficient was 0.941 **[Fig. 2C]**. In contrast, the correlation coefficient between the effects of untreated zymosan and depleted zymosan was only 0.518 **[Fig. 2C]**. Although zymosan and depleted zymosan exerted the same effect on some genes, such as *CXCL11, IFNB*, and *DDIT4*, they had disparate effects on other genes such as *SERPINB2, IL12B*, and *IL23*. Visualizing the correlation matrix for pairwise comparisons between all training regimens further strengthened the observation that ETSU β-glucan and depleted zymosan exert similar training effects while zymosan produces a distinct effect **[Fig. 2D]**. These analyses demonstrated that training with zymosan *vs*. β-glucans produces distinct transcriptomic responses to LPS.

We next sought to stringently identify genes that were differentially expressed after training. Among the 603 LPS-inducible genes, we identified 72 genes that were potentiated and 277 genes that were diminished by any training regimen, using a p-value cutoff of < 0.05 and a log2 fold-change cutoff (trained *vs*. untrained) of > 0.5. Of the diminished genes, 100 out of 277 (36.1%) were shared between untreated zymosan and the β-glucans **[Fig. 2E]**, consistent with the qualitative effect seen in Clusters 2, 3, and 4 of the heatmap **[Fig. 2A]**. In contrast, only 7 out of 72 potentiated genes (9.7%) were shared between untreated zymosan and the β-glucans, further evidence of their divergent training effects. The set of potentiated genes included key immune regulators such as *PTGIR* and *WNT5A* (potentiated by zymosan), *IL12B* (potentiated by β-glucans), and *BCAR1* (potentiated by both at the 3h timepoint) **[Fig. 2F**,**G]**. Given the small number of genes, no statistically significant enrichment of gene ontology terms was detected (using 603 LPS-inducible genes as the reference gene set).

### Zymosan and β-glucans activate distinct signaling networks

Having established differences between zymosan and β-glucans at both the secreted cytokine and transcriptomic levels, we hypothesized that structural differences between these ligands may lead to differential activation of cell signaling pathways to produce distinct training effects. We first performed proton nuclear magnetic resonance (NMR) to confirm the composition of each ligand. We found that untreated zymosan contained more lipid and protein, reflected in greater NMR signal between 0.5 and 2.8 ppm (24) **[Fig. S2]**. These NMR peaks were slightly diminished but still present in depleted zymosan, while the β-glucan from ETSU was nearly devoid of protein and lipid components.

All three ligands are known to engage Dectin-1 (32), but in murine myeloid cells and human cell lines untreated zymosan also engages TLRs, specifically TLR2 (21,22), in accordance with its higher protein and lipid content. Whether these ligand-receptor interactions hold true in primary human monocytes is uncertain. Furthermore, what nuclear transcription factors are activated downstream of these ligands in this system is not clear. To address these questions, we performed immunoblot on nuclear extracts of monocytes stimulated with zymosan or β-glucan. As ETSU β-glucan and depleted zymosan were shown to produce nearly identical training effects, we considered these two ligands as functionally equivalent and focused our signaling studies on ETSU β-glucan given its prominence in the literature.

We found that zymosan rapidly and robustly induced nuclear localization of NF𝓀B p65, phosphorylated AP-1 family member cJUN, and phosphorylated IRF3 **[Fig. 3A]**. Nuclear localization of STAT1 was also seen at later timepoints, likely due to secondary signaling from IRF3-induced type I interferon (IFN) (33). In contrast, ETSU β-glucan only weakly ac:vated NF𝓀B p65 and cJUN at 1 h, and did not detectably activate IRF3 or STAT1 **[Fig. 3A]**. Quantification of immunoblots showed that β-glucan-induced nuclear NF𝓀B p65 abundance was consistently less than zymosan-induced p65 at 1 h, with a mean reduction of 48% over four replicates, *p* = 0.020 **[Fig. 3B]**. β-glucan-induced nuclear accumulation of phosphorylated c-JUN was also less than zymosan-induced c-JUN at one hour, with a mean reduction of 74% over four replicates, *p* < 0.001 **[Fig. 3B]**. Taken together, these data indicate that zymosan activates signaling pathways that β-glucans do not, likely through engagement of additional signaling pathways.

**Figure 3:**
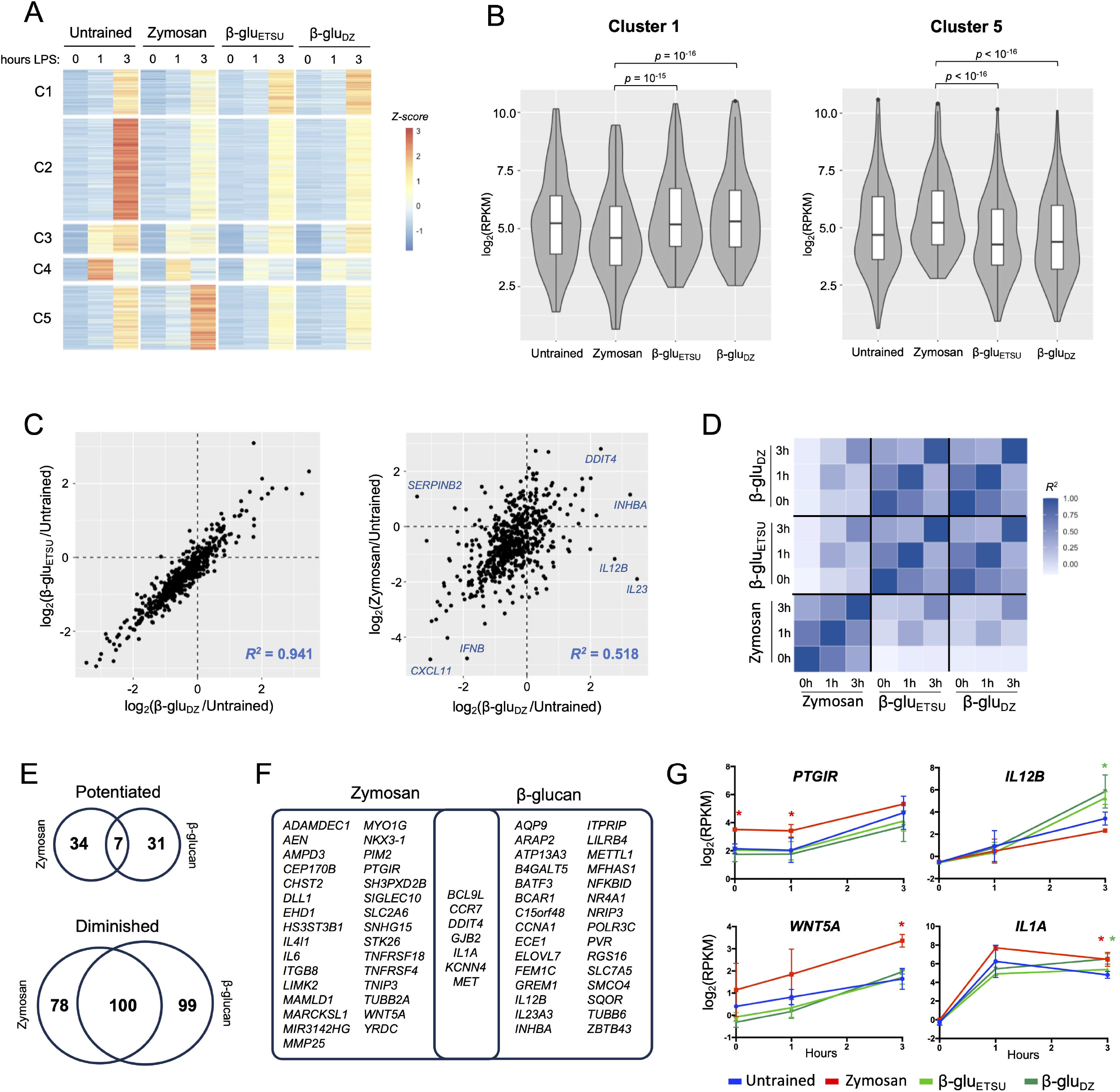
Zymosan and β-glucans activate distinct signaling networks. **(A)** Immunoblot of nuclear extracts of monocytes stimulated with zymosan or ETSU β-glucan over a 3-hour timecourse. Representative image from four replicates. **(B)** Quantification of nuclear NFκB (p65) and phospho-cJUN at 1 hour after stimulation with indicated ligands, normalized to p84, aggregate of four replicates.

### Concurrent exposure to TLR ligand, but not IFNβ, reverses β-glucan training

To test the hypothesis that differential activation of signaling networks produce distinct trained immunity phenotypes, we trained with mixtures of ligands using the same protocol as before, followed by secondary LPS stimulation and cytokine measurement. Pam3CSK4 is a TLR2-specific agonist that activates NF𝓀B and AP-1 pathways, while IFNβ activates STAT1 (33). We found that concurrent exposure to Pam3CSK4 reduced the β-glucan-mediated potentiation of TNF release by 51% (*p = 0*.*048*), while the addition of IFNβ had no effect **[Fig. 4A]**. These effects were largely conserved across a panel of cytokines measured by multiplexed bead array.

**Figure 4:**
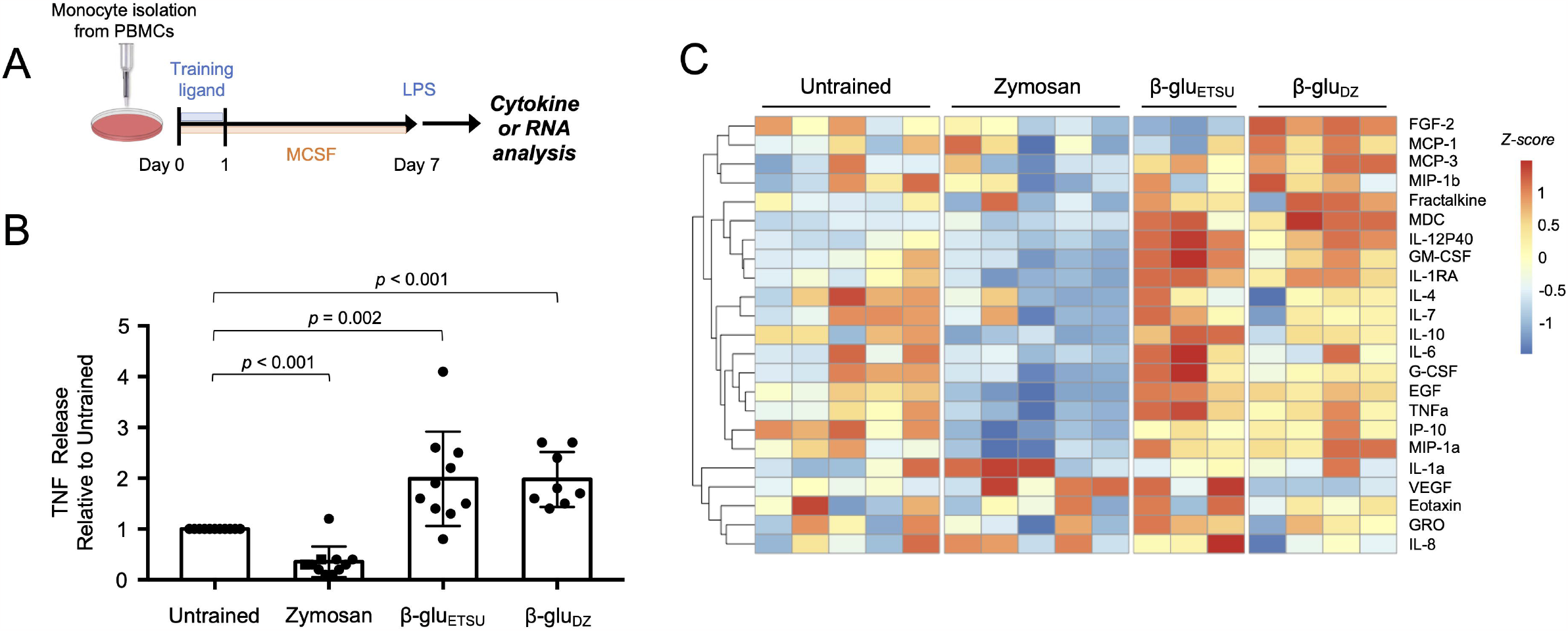
Concurrent exposure to TLR ligand, but not IFNβ, reverses β-glucan training. **(A)** TNF ELISA from supernatants of macrophages stimulated with LPS for eight hours, after training with the indicated ligands; p-values derived from two-tailed t-test, n.s. = not significant. Squares = ETSU, circles = depleted zymosan. **(B)** Heat map of multiplexed cytokine bead array from supernatant of macrophages stimulated with LPS for eight hours, after training with the indicated ligands.

Addition of Pam3CSK4 generally inhibited the training effects of β-glucan, producing a phenotype similar to zymosan, while addition of IFNβ preserved or in some cases enhanced the training effect of β-glucan **[Fig. 4B]**. Interestingly, for a small number of cytokines, the combination of β-glucan and Pam3CSK4 did not recapitulate the effect of zymosan. One of these exceptions was IL-1α, where zymosan potentiated cytokine release while β-glucan diminished it, and addition of Pam3CSK4 did not alter the β-glucan effect.

## Discussion

Innate immune training using prototypical stimuli such as β-glucan has potent effects, yet the determinants of s:mulus-specificity in trained immunity remain unclear. β-glucans train innate immune cells through the Dectin-1 receptor (12,16), but there are many different Dectin-1 ligands, both commercially available preparations as well as the widely published β-glucan from ETSU. These ligands have important differences in their preparation and structure that may lead to differences in innate immune training. We found that *C. albicans*-derived β-glucan from ETSU and *S. cerevisiae*-derived β-glucan (depleted zymosan) are nearly identical in their training effect. But unpurified zymosan induces a training regimen that is distinct from β-glucans, and in the case of many cytokines exerts the opposite training effect.

Our study clarifies that not all Dectin-1 ligands are equal with respect to innate immune training. It is particularly striking that alkali treatment of zymosan to reduce its TLR-activating properties produces a ligand (depleted zymosan) that has the opposite training effect on key cytokines such as TNF, IL6, IL12, and many others. It is also notable that one of the most widely used Dectin-1 agonists in the fungal pathogenesis literature (zymosan) produces dramatically different effects from the most widely published Dectin-1 agonist in the trained immunity literature (β-glucan from ETSU). Thus, investigators in both fields would be well-served to carefully consider their choice of Dectin-1 ligand.

One of the strengths of our study is the characterization of innate immune training across a wide array of secreted cytokines and the entire transcriptome. Much of the literature in this field has focused on archetypal inflammatory cytokines such as TNF, but relatively few studies have examined effects genome-wide. By utilizing systems-level analyses we show that innate immune training is both s:mulus-specific and gene-specific. That is, different stimuli produce different training effects, but each gene is uniquely regulated. For example, zymosan and β-glucans induce opposing training effects on IL-12, but they have the same effect on IL-1α. It is likely that studies which focus on one or two inflammatory cytokines underestimate the complexity of training effects. These studies have led to the proposal that “trained immunity” can be defined by a potentiation of inflammatory cytokines like TNF, while “tolerance” is defined by a reduction in the same cytokines (34). We suggest that rigid definitions of “training” and “tolerance” obscure subtle differences in immune training phenotypes, akin to the oversimplified paradigm of “M1” and “M2” macrophages (35).

At the signaling level, we found that zymosan activates the NF𝓀B and AP-1 transcription factors more strongly than β-glucan, and that co-activation of TLRs overrides the training program induced by β-glucan. While the activation of NF𝓀B, AP-1, and IRF3-STAT1 by zymosan corroborates what others have reported (32), the weak or non-existent activation of these key transcription factors by β-glucan was not anticipated. Remarkably, addition of the TLR2 ligand Pam3CSK4 largely reversed the training effect of β-glucans, whereas addition of IFNβ did not.

Others have shown that training with Pam3CSK4 and other TLR ligands diminishes TNF production in both murine and human macrophages (11,36). Our study extends these findings to demonstrate that when monocytes are concurrently trained with Pam3CSK4 and β-glucan, the TLR-induced activation of NF𝓀B and AP-1 pathways dominates sufficiently to inhibit the training effect of β-glucan.

Our observation that TLR activation opposes the training effect of β-glucans carries significant implications for the broader application of trained immunity. It is not known, for instance, how simultaneous exposure to different training ligands *in vivo* may alter response to subsequent infections. This is particularly relevant when considering that pathogens may train the immune system in one direction while simultaneously causing the release of cytokines and DAMPs that may produce opposing training effects. Thus, additional studies will be needed to define the mechanisms by which TLR activation opposes β-glucan-induced training. As trained immunity is thought to be encoded through modifications to the epigenome, one possibility is that TLR activation enforces an opposing epigenomic program. Stimulus-specific epigenomic reprogramming can occur even when similar transcription factors are activated due to differences in distinct patterns of temporal activity (37).

Interestingly, whole *C. albicans* yeasts, which activate both Dectin-1 and TLR2 (38), potentiate expression of TNF and IL6 (12). In fact, mannans sensed by TLR2 are required for maximal potentiation of TNF by *C. albicans in vivo* (16). While this seems at odds with our observations, three mechanisms could explain the apparent discrepancy. First, there are likely substantial differences in the relative availability of TLR2 binding sites between fragmented *S. cerevisiae*-derived cell walls (zymosan) and intact *C. albicans* yeasts (38). Secondly, it is possible that complex regulatory mechanisms exist *in vivo* that dampen the TLR-mediated effects (39); this would also explain why others have observed that zymosan injection *in vivo* slightly increases intracellular TNF in splenic macrophages (40). Thirdly, *C. albicans* interacts with multiple other host receptors *in vivo*, such as NLRP3, CR3, and mannose receptors, all of which have the potential to influence trained immunity (41).

Our understanding of the mechanisms underlying innate immune training remains incomplete, but this study advances the field by showing that across a broad range of cytokines and genes, Dectin-1 ligands produce distinct phenotypes of immune training. The fact that TLR activation reverses the training effect of β-glucan may provide important clues to further unravel the mechanisms of innate immune training.

## Author Contributions

QJC, KF, JF, ZM, SKM, and JS performed the experiments. QJC, KF, JF and ZM analyzed the data. QJC wrote the manuscript. KF and JF edited the manuscript, and all authors approved the final version. QJC conceived of and obtained funding for the study.

## Data Availability

Raw data and count tables for this study are available at Gene Expression Omnibus (GEO) under accession number GSE242947. Code used in data analysis can be provided upon request.

## Acknowledgements

This study was funded by NIH grant (to QJC) K08AI168567. It was performed in collaboration The Center for Genomics and Bioinformatics and the Immune Assessment core facilities at UCLA. We thank Alexander Hoffmann, Scott Filler, Alberto Yanez, and David Williams for their feedback and suggestions. We thank Hollie David for copyediting and administrative contributions.

