## Supplemental Figures for "Dectin-1 ligands produce distinct training phenotypes in human monocytes through differential activation of signaling networks"

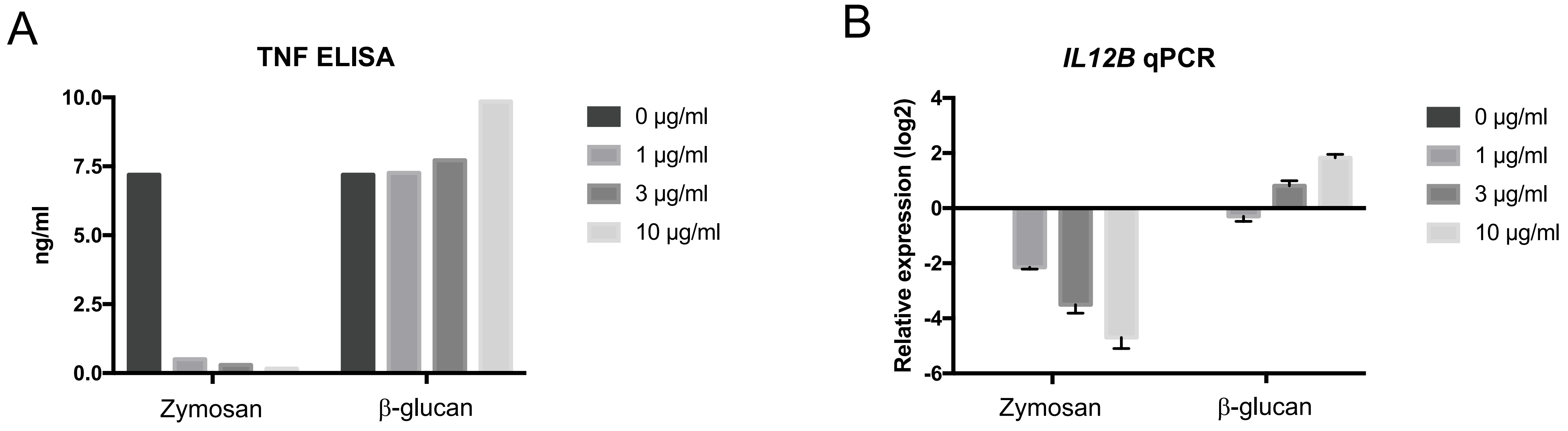


**Figure S1: Differential training effects are not a function of dose. (A)** TNF ELISA from supernatants of macrophages stimulated with LPS for eight hours, after training with the indicated doses of zymosan or β-glucan from ETSU. **(B)** qPCR for *IL12B* in macrophages, same conditions as (A).

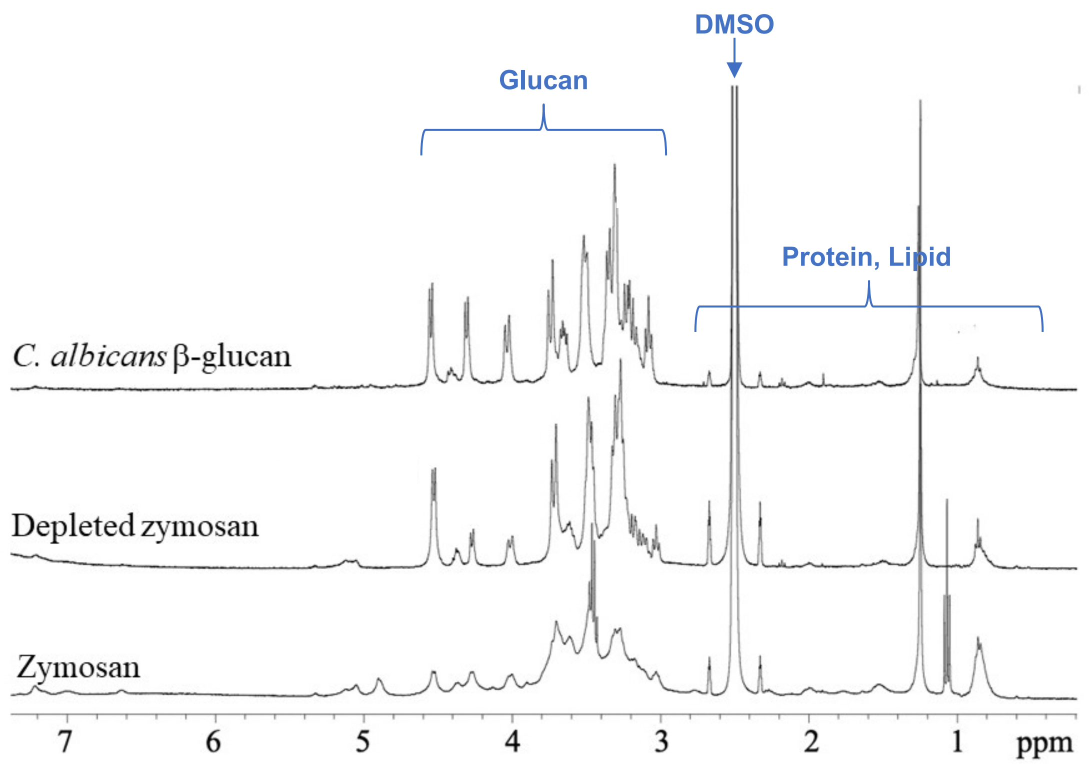


**Figure S2: NMR analysis of Dectin-1 ligands**
